# *millepattes* micropeptides are an ancient developmental switch required for embryonic patterning

**DOI:** 10.1101/376111

**Authors:** Suparna Ray, Miriam I Rosenberg, Hélène Chanut-Delalande, Amelie Decaras, Barbara Schwertner, William Toubiana, Tzach Auman, Irene Schnellhammer, Matthias Teuscher, Abderrahman Khila, Martin Klingler, François Payre

## Abstract

Small open reading frames (smORFs) that code for “micropeptides” (10-100 amino acids) exhibit remarkable evolutionary complexity. Conserved micropeptides encoded by the *millepattes (mlpt)* gene are essential in *Tribolium* for embryogenesis but in *Drosophila,* function only in leg and cuticle differentiation. We find that a module identified in *Drosophila* trichome patterning, comprising Mlpt, UBR3, and Shaven-baby (Svb), coordinates early embryo patterning in several insect orders. Intriguingly, Mlpt segmentation function can be re-awakened in the *Drosophila* blastoderm, demonstrating the potency of an ancestral developmental switch retained despite evolving embryonic patterning modes. smORFs like *millepattes* thus illustrate plasticity of micropeptide functions despite constraints of essential genetic networks.

**One sentence summary:** A module comprising the small ORFs mlpt/pri/tal, the transcription factor Svb, and the ubiquitin ligase UBR3, possesses an ancestral function in insect embryo patterning which is lost in flies but reactivated when Svb expression is restored.

## Introduction

Small open reading frames (or smORFs) are abundant in all genomes, but few have been investigated. One of the first smORFs to be deeply studied phenocopies the unusual phenotype of Shaven-baby (Svb), a transcription factor that is a central node in epidermal patterning in flies (Payre F et al, 1999). Larvae lacking *svb* are left without epidermal trichomes, due to failure to activate the actin remodeling program underlying their differentiation. The smORFs encoded by the insect-specific *millepattes* (*mlpt)* locus have now been described and studied in several insect developmental systems. The peptides are deeply conserved throughout insects, a taxon representing over 300 million years of evolution. Two contemporary studies independently identified this gene in *Drosophila*, naming it “*tarsalless”* (*tal*) as a regulator of proximal-distal leg patterning (Payre F et al 1999, Pueyo JI and Couso JP.,2008) and “*polished rice”* (*pri*) for its essential role in formation of trichomes in the larval cuticle (Galindo MI, et al 2007). In the epidermis, Pri peptides enable recruitment of the proteasome via the E3 ubiquitin ligase, UBR3, to the transcription factor shaven-baby. Svb, a transcriptional repressor, is converted to a transcriptional activator through proteasome-mediated cleavage and degradation of its N-terminal repression domain (Zanet J. et al 2015). The locus encoding Pri peptides exhibits detectable synteny and encodes conserved peptides of varying number throughout the insect lineage (Galindo MI, et al 2007, Kondo T, et al 2007, Savard J et al 2006). Independently, Savard et al. (2006) discovered an essential function for this locus in the formation of abdominal segments in the so-called “short-germ” flour beetle, *Tribolium castaneum*. In beetles, RNAi knockdown of *mlpt* caused loss of abdominal segments (posterior truncation of the embryo), as well as transformation of anterior abdominal segments to thoracic fate, leading to a distinctive phenotype of extra pairs of legs (*millepattes* is French for centipede). Strikingly, no early embryonic patterning phenotype was observed for *mlpt* (*tal, pri*) in *Drosophila*, in spite of its expression pattern, which resembles the striped expression of *Drosophila* hairy, a critical segmentation gene in flies (Galindo MI, et al 2007, Kondo T, et al 2007; Figure 5). This apparent paradox left open the question of how Mlpt peptides may function in segmentation in *Tribolium.*

## Results

### Identification of mlpt partners Svb and Ubr3 in Tribolium segmentation

In the genome-wide iBeetle RNAi screen in *Tribolium* (Schmitt-Engel, C. et al 2015) we identified *ubr3* on the basis of its cuticle phenotype that strongly resembles that of *mlpt* (Table S1; Fig 1B,C), which suggested that the complete fly module comprising *mlpt/svb/ubr3* may be conserved. Indeed, targeted knockdown of *svb* by parental RNAi leads to an abdominal truncation and homeotic transformation phenotype that closely resembles that of *mlpt* and *ubr3* knockdowns (Figure 1 D, E; Savard J et al 2006; Materials and methods). In addition to segmentation defects, these phenotypes share several similarities, including transformation of abdominal segments towards thoracic identity, shortened sensory bristles, shortened leg segments with a “bubble-like” terminus, and missing telson appendages. (Fig 1. B-G). Perturbed animals also exhibit subtle phenotypic differences in number of ectopic leg bearing segments and body length. Severe *ubr3* RNAi phenotypes are stronger, often with severely shortened larvae missing most abdominal segments, while embryo RNAi even at very low concentrations is lethal (no cuticles develop), supporting the involvement of this ubiquitin E3 ligase in *mlpt*-independent functions. ubr3 RNAi leg segments are also more rounded and the pretarsi shorter. Knockdown svb larvae are characterized by the presence of legs on the first two abdominal segments only; one or more legs on segment A1 are often reduced. Presence of T2-like spiracles on A1 and the absence of spiracles on A2 in *svb* knockdowns suggest transformation of the two anterior abdominal segments into thoracic segments, T2 and T3, respectively. Leg segments are more severely shortened and rounded and pretarsi more strongly reduced in svb knockdowns than in mlpt knockdowns (Fig 1, Supplement).

CRISPR knock-out of *svb* highlights an additional phenotype not apparent from RNAi knockdown, which is considerable thinning of the epidermal cuticle similar to what has been observed in the fly (Andrew DJ and Baker BS, 2008; Figure 4; Figure 1, Supplement 3). These results strongly support similar functions of Mlpt, Ubr3 and Svb in *Tribolium* and flies. Moreover, similarities in transcript, and protein structure and disorder disposition profiles between *Drosophila* and *Tribolium* Svb suggest similar molecular interactions (Figure 1-supplement 1). As in flies, we observe that *Tc-mlpt* knockdown embryos accumulate the repressor form of Svb protein, as detected by an antibody recognizing the N terminal repression domain of *Drosophila* Svb (Figure 1-supplement 1). Taken together, our data support a cooperation among Svb, Ubr3 and Mlpt peptides, defining a conserved module essential for *Tribolium* segmentation.

**Figure 1.**
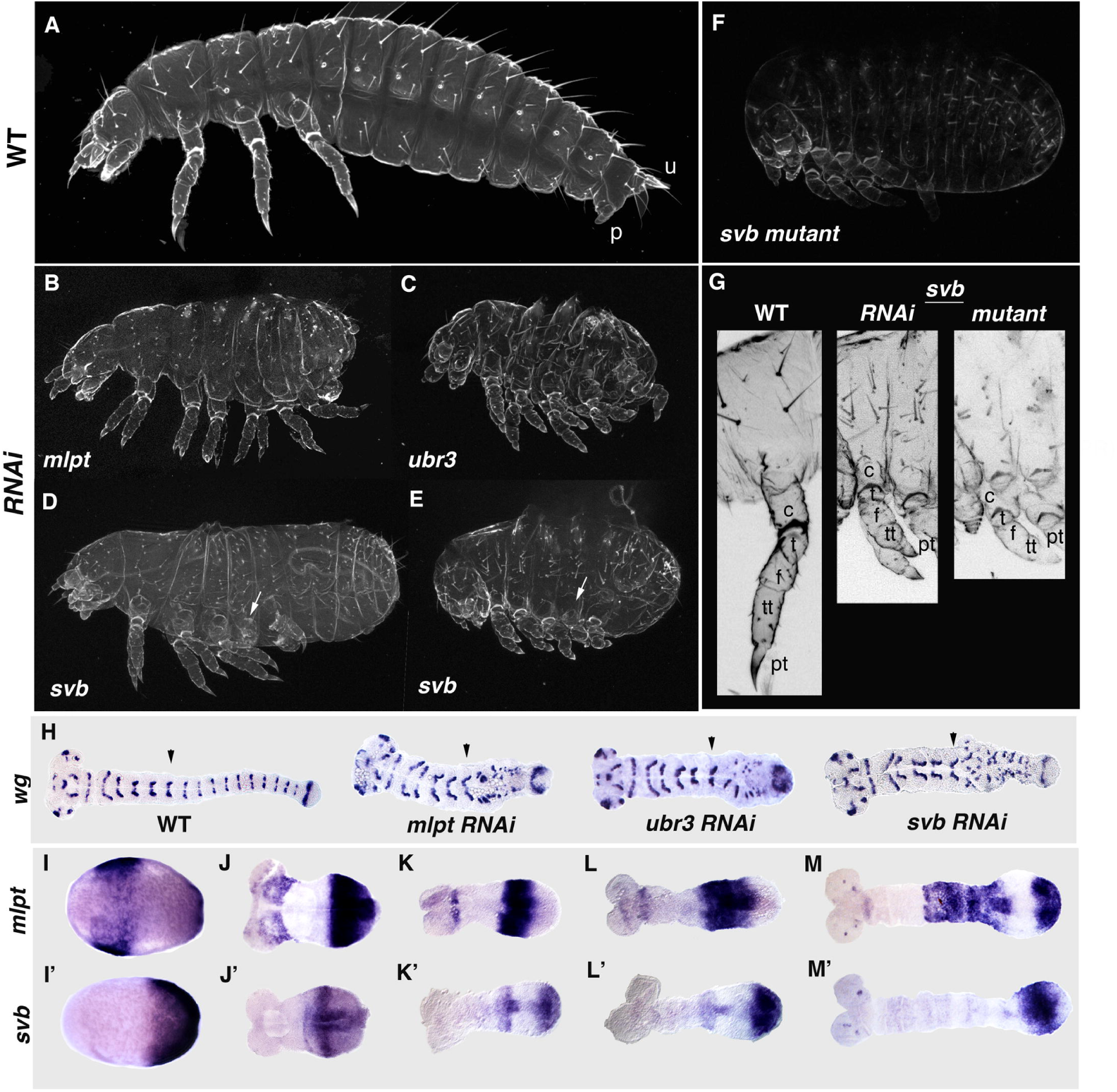
Loss of function embryonic phenotypes of *mlpt, ubr3*, and *svb* and their embryonic expression reveal a cooperative segmentation function of the module in *Tribolium castaneum (Tribolium)*. (A-F) Cuticle phenotypes of first instar larvae of *Tribolium* WT [A], *mlpt* [B], *ubr3* [C], *svb* [D, E] RNAi, and a knock-in *svb* CRISPR mutant [F]. *mlpt, svb*, and *ubr3* knockdown larvae after parental RNAi are shortened along the anteroposterior axis and are more compact relative to wild type. While only the first two abdominal segments are transformed in *svb* knockdowns, almost all abdominal segments can be transformed to thoracic identity in *mlpt* and *ubr3* knockdowns. One or both legs on the transformed A1 segment are usually reduced to stumps in *svb* knockdowns (white arrows; D, E). D and E portray weaker and stronger *svb* RNAi phenotypes respectively. To create the *svb* CRISPR mutant, guide RNAs targeting the transactivation domain of *svb* were used to knock-in a plasmid bearing EGFP under the control of a 3xP3 promoter into the *svb* locus by inducing double-stranded breaks followed by non-homologous end joining (NHEJ). *svb* mutant cuticles have very low autofluorescence and appear dim [F]. WT, wild-type; u, urogomphi; p, pygopods. (G) Comparison of the prothoracic leg from a *Tribolium* WT, a *svb* RNAi, and a *svb* mutant larva. Leg segments of the *svb* knockdown and mutant larvae are significantly shortened and rounded compared to the WT and the pretarsus, significantly reduced. Bristles are missing on the mutant leg and the pretarsus is not sclerotized. coxa (c); trochanter (t); femur (f); tibiotarsus (tt); pretarsus (pt) (H) *wingless* expression in *Tribolium* WT, *mlpt, svb*, and *ubr3* RNAi embryos. The stripe corresponding to the 3^rd^ thoracic segment (T3) is shown (black arrows). In all three knockdowns, segmental stripes can be disrupted right after the T3 stripe has formed. (I-M’) Whole mount *in-situ* hybridization results showing the expression of *Tc-mlpt* and *Tc-svb* from late blastoderm through extending germband stages highlighting their complementarity of expression. In the late blastoderm *mlpt* and *svb* expression domains overlap in the posterior (I and I’). In the late germ rudiment *svb* is strongly expressed in an anterior band with diffuse expression in the posterior; *mlpt* is strongly expressed in the diffuse *svb* domain (J and J’). In the early germband, *svb* expression clears out in the middle forming an anterior and a posterior band; intense *mlpt* expression is present in the middle region with sparse *svb* and absent at the posterior end (K and K’). As the germband grows, the anterior domain of *svb* fades while its posterior domain remains strong abutting on the posterior border of the *mlpt* 1^st^ trunk domain (L and L’). The 1^st^ trunk domain of *mlpt* fades while a 2^nd^ one is established at the posterior; *svb* domain detaches from the posterior and is concentrated in a region close to the posterior end, roughly between the two *mlpt* trunk domains (M and M’).

Notably, expression of the segmental marker *wg* confirms that only abdominal segments are disrupted in *mlpt, svb*, and *ubr3* RNAi embryos while thoracic segments are formed normally (Fig. 1 H). This is of interest since in the “short germ” embryo of *Tribolium* only head and thorax form in the “syncytial” or acellular environment of dividing nuclei in the blastoderm, while after cellularization, the embryo undergoes convergent extension, and abdominal segments continue to arise in a sequential and regular manner from the posterior “growth zone” (Rosenberg MI et al, 2009, Liu PZ and Kaufman TC, 2005). In *Drosophila*, although both *pri/mlpt* and *svb* are expressed throughout embryogenesis, there is little overlap in their expression domains until after germband retraction, as *pri/mlpt* is expressed in stripes resembling pair rule stripes, while *svb* is expressed only in the head (Figure 4); *ubr3* expression is ubiquitous. Since functional conversion to an activator via cleavage of Svb protein relies on the presence of Pri/Mlpt peptides, the timing and co-incidence of their expression is important. The observation that loss of *pri* segmentation function in flies coincides with absence of *svb* expression in the early fly embryo led us to investigate the mRNA expression of Mlpt partners Svb and Ubr3 in *Tribolium.*

We find that, as in the fly, *svb* and *mlpt* mRNA are expressed during both blastoderm and germband stages of embryogenesis in *Tribolium,* and importantly, are co-expressed in the early embryo as *Tribolium svb* mRNA is expressed maternally (data not shown). Zygotic expression of *Tribolium svb* also overlaps with *mlpt* in the pre-growth zone at the onset of gastrulation (Fig. 1I, 1I’). In the extending embryo, this overlap resolves into complementary expression: the posterior *svb* domain evolves into a strong anterior band and a more diffuse posterior expression (Figure 1J’’) while *mlpt* has much stronger posterior expression (Figure 1J). As the germ rudiment matures, *svb* forms two distinct expression domains flanking the *mlpt* expression domain (Figure 1K, K’). High levels of *mlpt* and *svb* expression may be mutually repressive (Figure 1, supplement 5). Subsequently, *svb* and *mlpt* expression domains shift, wave-like, anteriorly while anterior *svb* expression fades and its posterior expression detaches from the posterior end (Fig. 1M’). Concomitantly, *mlpt* trunk expression moves anteriorly and a second posterior domain arises that is again complementary to the now anteriorly shifted *svb* domain (Fig. 1 M), with fading anterior *mlpt* overlapping with the fading anterior *svb* domain (Fig. 1M and M’). The interaction at the interfaces of the complementary domains may be critical for patterning of the abdominal segments. Posterior trunk expression of *mlpt* eventually also detaches from the growth zone and gradually fades while *svb* persists as a strong narrow band of expression until segmentation is completed (Figure 1-supplement 2). Late *svb* is expressed in presumptive neurons during germ band extension, and later, in the head and appendages (Figure 1M’, Figure 1-supplement 2).

In summary, *Tribolium mlpt* and *svb* expression patterns are dynamic and often complementary, though at times, overlapping. Loss of function phenotypes of *Tc mlpt, svb* and *ubr3* suggest that a functional module for *mlpt* discovered in *Drosophila* trichome patterning also works in concert in embryonic segmentation, leg patterning and cuticle formation in *Tribolium.*

### Complementarity of expression of pri/mlpt and svb is conserved deeply in insects

Our data revealed a surprising and essential role for this gene module in controlling posterior segment formation and identity in *Tribolium.* To determine whether these genes are similarly important for embryonic development of other insects, we investigated their expression patterns in additional, more basal insect species: the water strider, *Gerris buenoi* (“Gb”; Hemiptera, Gerridae) and the milkweed bug, *Oncopeltus fasciatus* (“Ofas”; Hemiptera, Lygaeidae). Figure 2 highlights the expression patterns of these genes at several timepoints in embryogenesis. The early development of water strider and milkweed bug are quite similar. *Ubr3* expression is ubiquitous in both *Tribolium* and *Gerris* and was not examined further. *mlpt* and *svb* expression in the early hemipteran embryo is observed in adjacent domains at the anterior of the precellular blastoderm embryo (e.g. *Oncopeltus,* Figure 2A-A’), with additional strong posterior *svb* expression at the future site of invagination which becomes broad expression throughout the early growth zone (Figure 2A; Figure 2-supplement 1). This complementarity perdures, until a transition to transient overlap in the early growth zone (Figure 2-supplement 1D-D’). Subsequently, expression domains of *svb* and *mlpt* again resolve into complementary “waves” of *svb* and *mlpt* stripes moving anteriorly within the growth zone (Figure 2B-B’; 2D-E’; Figure 2-supplement 2); *Ofas mlpt* expression is also diffusely expressed through recently added segments anterior to the GZ (Figure 2C’). Later expression in both species is seen in the limb buds and mouth parts (Figure 2C-C’, E-E’), consistent with a function in patterning the leg and head appendages, as well as in presumptive neurons of the head and along the midline. These data hint at a surprising role for this gene module in controlling segment formation and identity in representatives of the Coleoptera and Hemiptera, but not Diptera.

### Conserved function of mlpt gene module in insect segmentation

We next tested whether and how broadly *mlpt, svb*, and *ubr3* may functionally cooperate during embryogenesis in these additional short germ insects. We used RNAi against each of these genes in water striders and milkweed bugs, and find that all cause severe patterning defects.

**Figure 2.**
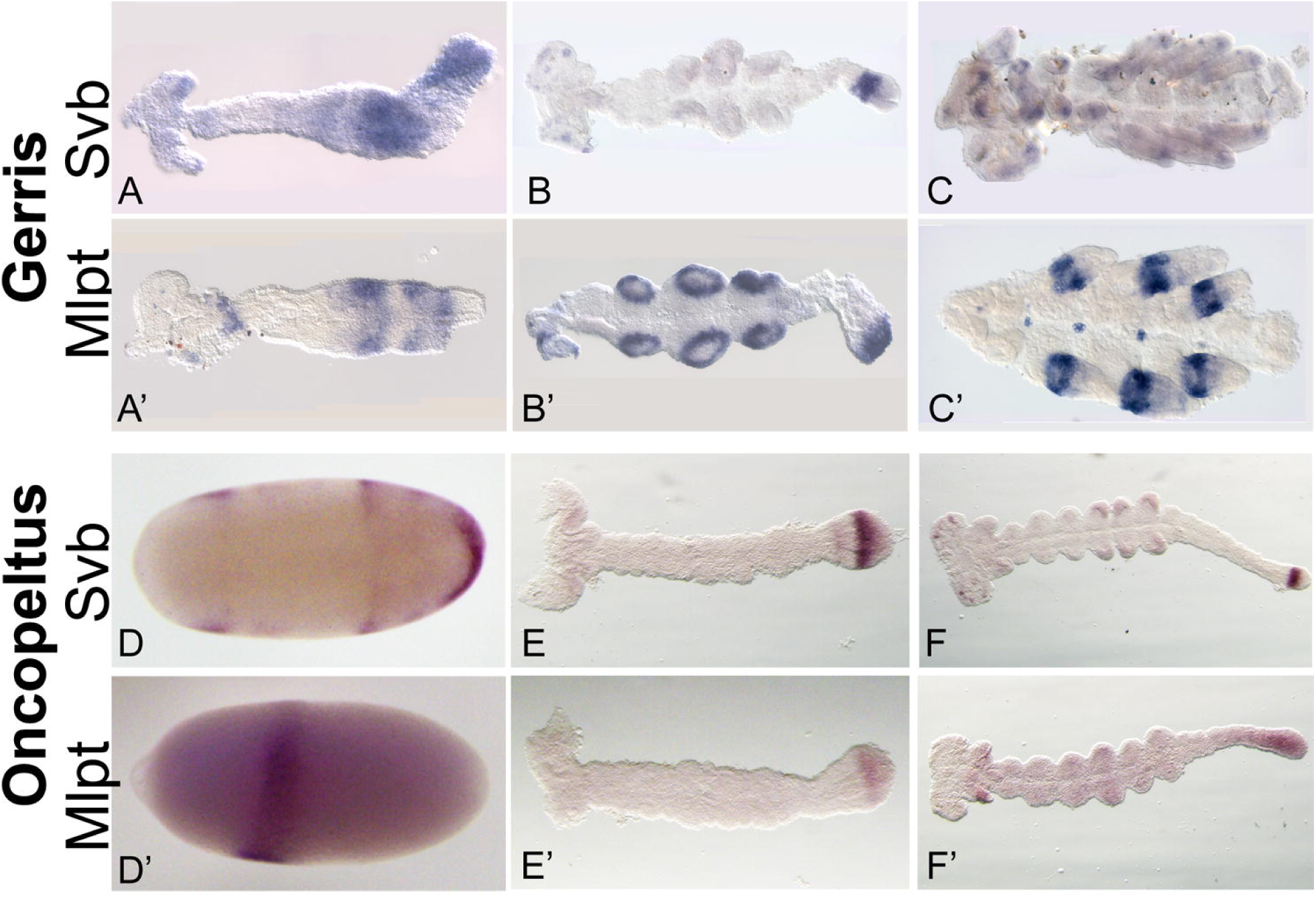
Embryonic expression of mlpt and Svb mRNA in hemipteran embryos of *Gerris buenoi* and *Oncopeltus fasciatus.* (A-C) Expression of *svb* in the *G. buenoi* embryo. (A) *G.bue Svb* expression is evident as faint expression the head and thorax, along with stronger domains of expression in the abdomen of the early germband. (B) Mid germbands exhibit more restricted expression of *Svb*, in a stribpe in the growth zone, as well as in putative neurons in the head, and faint expression in the limb buds. (C) Late stage embryos exhibit banded expression of *Svb* in the legs and head appendages, as well as in foci of expression in the head. **(A’-C’)** Expression of *mlpt* in the *G. buenoi* embryo. (A’) *G.bue mlpt* expression in the early germband is restricted to a thoracic stripe and two distinct domains of abdominal expression, which abut domains of *G.bue Svb* expression (A). (B’) Mid germband embryos exhibit strong staining in the limb buds, as well as in the extreme posterior of the growth zone, immediately adjacent to a strong domain of *G. bue Svb* in the middle of the growth zone (B’). (C’) Late stage *G. bue* embryos exhibit strong staining in the mature limbs, as well as in foci of expression along the midline of the embryo. **(D-F) mRNA expression of Svb in the *Oncopeltus fasciatus* embryo**. (D) A dorsal view of a blastoderm embryo of *Oncopeltus* highlights its two domains of expression: one anterior domain of head expression, and a posterior domain encompassing thoracic segments, and strong staining at the site of ingression where gastrulation begins. (E) Early germband staining for *Ofas Svb* transcripts reveals faint staining in the headlobes, as well as strong staning in two distinct stripes in the growth zone. (F) Later germbands express *svb* in putative neurons of the head, as well as in limb buds and a strong stripe in the middle of the growth zone. (**D’-F’) mRNA expression of *Ofas mlpt* in the Oncopeltus embryo**. (D’) Blastoderm expression of *Ofas mlpt* is slightly less robust than *Svb* expression, and is restricted to a single strong strip corresponding to the future head segments. (E’). Early germband embryos exhibit a single domain of *Ofas mlpt* expression in the extreme posterior of the growth zone, posterior to the strong mid-growth zone domain of *Ofas Svb* expression. (F’) Mid-germband *Ofas* embryos exhibit strong expression in the head appendages, as well as broad low level expression in the putative head and thoracic segments and strong but diffuse expression throughout the growth zone.

Embryos of hemimetabolous insects, including water striders and milkweed bugs, complete embryogenesis and undergo a series of molts (which does not include a pupal stage) through which these hatchlings eventually achieve adult size and morphology. These intermediate nymph stages exhibit the full complexity of adult structures, allowing for detailed phenotypic analysis. In water strider and milkweed bugs, the wildtype hatchling possesses three long pairs of legs, which extend along the ventral side, curling around the posterior, as well as a long pair of antennae that extend posteriorly along the ventral midline (Figure 3A-B; 3I-J). *mlpt* RNAi in both *Gb* and *Ofas* results in loss of posterior abdominal segments and fusion of thoracic segments, with shortened rounded legs that terminate proximal to the trunk; reduction and fusion of head appendages is also apparent (Fig 3C-D, 3K-L). In *Ofas*, severely affected embryos fail to gastrulate, resulting in an everted gut. *Gb* and *Ofas svb* RNAi also results in loss of abdominal segments and rounding of more distally truncated legs (Figure 3E-F, 3M-N). *ubr3* RNAi in both species gives the most severe phenotype, reflecting its presumed additional functions independent of *svb* and *mlpt. Gb ubr3* RNAi embryos exhibit loss of posterior abdomen, as well as fusion or loss of most leg-bearing segments and reduced head appendages and eyes; remaining legs are fused and shortened with bubble-like termini (Figure 3G-H). In *Ofas*, severe *ubr3* RNAi embryos are almost completely ablated, leaving unidentifiable ectodermal tissue connected to everted presumptive visceral tissue. More mildly affected embryos possess some apparent segment identity, with head and eyes, but no appendages and limited evidence for correct axial polarity (Figure 3O-P).

**Figure 3.**
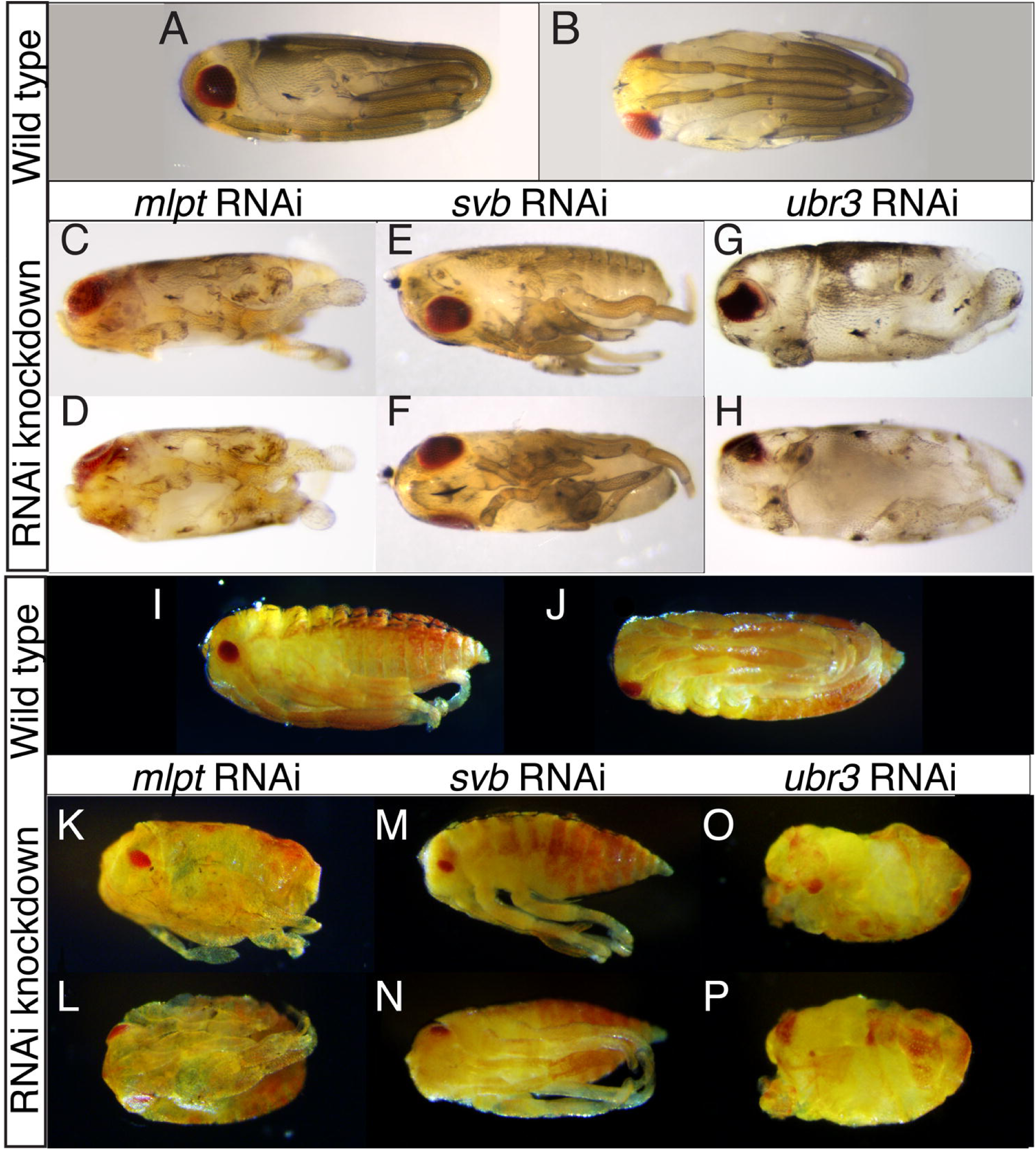
Knockdown of *mlpt, svb*, and *ubr3* function by RNAi in Hemipteran embryos results in a shared subset of segmentation and patterning phenotypes. (A-H)Gerris buenoi phenotypic study of *mlpt, svb, and ubr3* RNAi. Wild type Gerris buenoi hatchlings from a (A) lateral and (B) ventral perspective. *Gbue* hatchlings possess visible bristles on a pigmented epidermis, red pigmented eyes of characteristic shape and size, and elongated mouth parts and antennae that extend along the ventral side of the embryo, terminating between the long legs which wrap around the posterior end of the embryo. (C-D) *Gbue mlpt* RNAi embryos (shown laterally in C, ventral view in D) exhibit posterior truncation, resulting from loss of posterior abdominal segments, as well as loss and/or fusion of legs and head appendages and alterations in eye morphology. (E-F) *Gbue Svb* RNAi embryos (show laterally in E, ventral view in F) also exhibit posterior truncation with shortening or fusion of legs and head appendages. (G-H) *Gbue ubr3* RNAi embryos (shown laterally in G, ventral view in H) exhibit more severe phenotypes, which include posterior truncation and more extreme alterations/loss of legs and head appendages, as well as more severe disruption of the eye. **(I-P)** Embryonic phenotypes of *Oncopeltus fasciatus* (*Ofas*) *mlpt, svb*, and *ubr3* RNAi. Wild type *Ofas* hatchlings from a lateral (I) and ventral (J) perspective, showing typical pigmentation, and typical characteristic size, shape and number of appendages. (K-L) Parental RNAi of *Ofas mlpt* causes severe posterior truncation, loss of abdominal segments, and fusion/loss of thoracic segments, and shortened legs and head appendages and a reduced eye. (M-N) Parental RNAi of *Ofas svb* causes fusion/loss of thoracic segments and reduction of both legs and eyes. More severe RNAi phenotypes include posterior truncation of the embryo, loss/fusion of legs and head appendages. (O-P) Parental RNAi of *Ofas ubr3* causes severe truncation of the embryo, loss of all appendages and reduction of the eye, as well as apparent loss of axial polarity/axis duplication in severely affected embryos. In addition, severely affected embryos of all genotypes exhibited significant thinning of the cuticle.

### Conservation in alternative long germ insects

Since all basal insect species examined showed evidence of conserved function of this module in segmentation, we tested whether other more derived species may retain this functionality. Like *Drosophila, Nasonia, Apis*, and other Hymenoptera exhibit “long germ” embryogenesis, in which the embryo is mostly patterned in the context of the syncytical blastoderm, and which has evolved several times in the insect phylum (Rosenberg MI, et al 2009, Liu PZ and Kaufman TC, 2005,Misof B et al 2014). *Drosophila svb* expression in the early embryo is restricted to two stripes in the head (Mével-Ninio M et al 1995, Figure 4 A,B), while *Dmel pri* is expressed throughout the blastoderm, resembling the striped expression of the pair-rule gene *hairy* (*3*). In contrast, in the long germ embryo of *Nasonia vitripennis, mlpt* and *svb* are expressed in the precellular blastoderm in adjacent posterior domains (Figure 4E-F’), at the region of the wasp embryo described previously as the progenitor of the late-forming segments (Rosenberg MI et al 2014). *Nv svb* is also expressed in a prominent stripe in the middle of the embryo, similar to expression of the thoracic gap gene, *Krüppel* (Brent AE et al 2007; Figure 4E), while *mlpt* expression has an anterior cap, and broad expression posterior to the *svb* domain (Figure 4E’-F’). Later expression of both genes in flies can be seen in segmental stripes during germband extension that prefigure the larval trichomes (Figure 4C-D’), while in *Nasonia,* both genes have expression on the dorsal side of the embryo, beginning around gastrulation (Figure 4G’; Figure 2-supplement 2). Late expression of *svb* in *Nasonia* appears in stripes, prefiguring the larval trichomes. Late expression of *mlpt* is also in stripes, which are reminiscent of segmental genes *en* and *wg*, and coincident with *svb* expression, as well as in spots in the presumptive nervous system (Figure 2-supplement 2). Thus, in a wide range of insects, complementary and sporadic overlapping expression of *svb* and *mlpt* in the embryo correlates with an essential role in embryonic segmentation.

In holometabolous insects, like *Drosophila* and *Nasonia*, the embryo hatches into a larva, whose cuticle is ornamented with dense bristles called trichomes. Since the pattern of trichomes, or denticles, is predictable and distinctive along the anterior-posterior (A-P) and dorsoventral axes and within each segment, it can serve as a readout for correct segmentation. The fly larva has a characteristic trichome pattern in WT flies (Figure 4I), which is severely reduced (hence, “shaven”) in the thin cuticles of larvae mutant for *svb, pri,* or *ubr3* (Figure 4J-L). However, all segments are still formed in these mutants. The larval cuticle pattern of *Nasonia* differs from the fly; three thoracic and 10 abdominal segmental bristle bands can be discerned, as well as 4 spiracles, located on thoracic segment T2, and abdominal segments A1, A2 and A3, which provide essential landmarks for segment identification (Pultz MA et al 2000; Figure 4M). In *Nasonia, mlpt* RNAi causes posterior truncation and segment fusions, evident as a severely shortened larval cuticle, with two remaining bands of denticles that are likely thoracic and anterior abdomen (Figure 4N). Similarly, *svb* RNAi causes severe posterior truncation and loss of most abdominal segments. Two clear bands of denticles and two pairs of spiracles remain (Figure 4O). Larvae from *ubr3* RNAi were almost uniformly too fragile to recover (not shown), likely owing to the observed absence/thinning of cuticle; mildly affected *ubr3* RNAi larvae exhibit thin cuticle, devoid of denticles on the posterior, with anterior fusion of remaining denticles (Figure 4P). Altogether, our data support conserved functions for *mlpt, svb* and *ubr3* in embryonic segmentation of a long germ insect, leaving only long germ *Drosophila* from among species tested without such an early patterning function.

**Figure 4.**
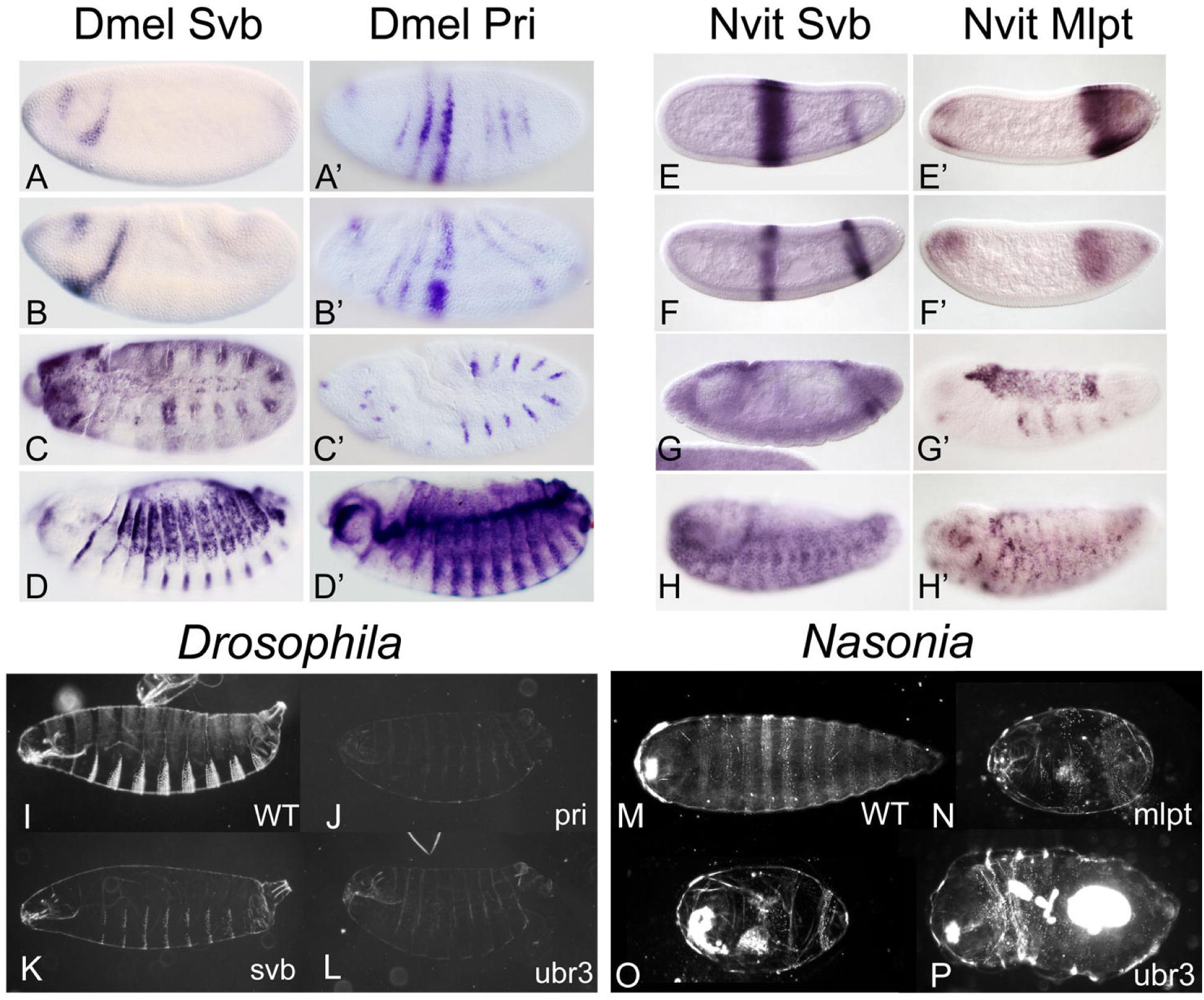
Expression and loss of function phenotypes of *svb,* and *mlpt*/*pri* in the long germ embryos of *Drosophila melanogaster* and *Nasonia vitripennis*.(A-D’) Expression of *Dmel svb* and *pri* mRNA in the *Drosophila* embryo. (A-D) *D. melanogaster* embryonic *Svb* mRNA expression. (A) *D.melanogaster* embryos express Svb mRNA in the pre-cellular blastoderm and (B) gastrula embryo, where it is expressed in two stripes in the head. (C) Embryonic expression of *D.mel Svb* at germband extension. Expression at this stage is more widespread, including stronger staining in a segmental pattern of stripes. (D) Dorsal closure embryo expression of *D.mel Svb* mRNA. At the final stages of embryogenesis, strong expression of *Dmel svb* is seen in a pattern that prefigures the denticles, apical projections of actin that form small bristles on the larval cuticle and which have a stereotypical pattern on both the dorsal and ventral sides of the *Drosophila* larva. (A’-D’) *D.melanogaster* embryonic *pri* mRNA expression. (A’) Precellular blastoderm expression of *D.mel pri* mRNA. *pri* mRNA is expressed in seven thin, apparently parasegmental stripes plus a head stripe (similar to the blastoderm expression of pair-rule gene, *hairy*), of which all but one stripe perdure through gastrulation (B’). (C) Germband extension embryo expression of *D.mel pri* mRNA. Ten segmental stripes of *Dmel pri* expression are evident at full germband extension, with additional expression in presumptive neurons in the head. (D) Strong *Dmel pri* expression is observed throughout the dorsal closure embryo, with the darkest staining highlighting the trachea and prefiguring patterns of dorsal and ventral larval denticles, coincident with *Dmel Svb* mRNA expression. **(E-H’)** Expression of *Svb* and *mlpt* mRNA in the *Nasonia vitripennis* (*N.vit*) embryo. *N.vit Svb* is expressed in the mid (E) and late (F) blastoderm embryo in two stripes. The anterior stripe, initially more robust, is in the middle of the embryo, resembling the expression of gap gene, *N.vit Krüppel (ref Ava)*. A more posterior stripe of *N.vit Svb* is expressed in the same region of the embryo where the most posterior stripe of *N.vit eve* is expressed, and later gives rise to 6 additional posterior segments. These embryos also exhibit a strong stripe of dorsal expression. (G) The gastrula embryo of *Nvit* exhibits low level *Svb* expression throughout the embryo, with some enrichment at the cephalic furrow; the posterior stripe of *N.vit Svb* splits into two stripes at gastrulation and the central thoracic stripe is no longer seen. Dorsal expression of *Nvit Svb* is also lost during gastrulation. (H) By dorsal closure, *Nvit* embryos exhibit widespread *Svb* mRNA expression, with visible enrichment in a segmental pattern and prefiguring the pattern of larval denticles, as in *Drosophila.* (E’-H’) Embryonic expression of *N.vit mlpt* mRNA. (E’) Mid blastoderm expression of *N.vit mlpt* mRNA. *Nv mlpt* RNA is expressed in an anterior cap and a stronger posterior domain, covering the entire posterior of the embryo, but enriched in a strong stripe at the anterior of this domain. (F’) Late blastoderm expression of *N.vit mlpt* RNA includes a head domain of similar intensity, is weaker, and also withdrawn anteriorly from the embryo posterior. This leaves a broad posterior stripe overlaying the region of posterior *N.vit eve* expression, and a small domain of more intense expression underlying the pole cells. (G’) Germband retraction embryo expression of *N.vit mlpt*. After gastrulation, during which *mlpt* expression is observed on the dorsal side of the embryo, and in two stripes (at the middle and at the posterior of the embryo), dorsal expression is enhanced as the embryo approaches dorsal closure, with the addition of several stripes of expression along the lateral ectoderm. (H’) Late embryo expression of *N.vit mlpt*. At the end of embryogenesis, *N.vit mlpt* expression is observed in presumptive neurons in the head and throughout the embryo, in stripes of ectoderm in each segment, as well as in additional cell types throughout the head and trunk of the embryo. **(I-L)** Cuticle preps of *Drosophila melanogaster* larvae. (I) Cuticle of wild type *D. mel* larva showing typical pattern of ventral denticles. (J) Cuticle of *D.mel pri* mutant larva, showing both cuticle defects and absence of denticles. (K) Cuticle of *D.mel Svb* mutant larva, showing severe depletion of ventral denticles, as well as thinning of larval cuticle. (L) Cuticle of *D.mel ubr3* mutant larva, showing absence of denticles and thinning cuticle. **(M-P)** Cuticle preps of *Nasonia vitripennis* larvae. (M) Cuticle of wild type *N.vit* larva, showing stereotypical pattern of denticles, and spiracles that serve as positional identity landmarks, on segments T2, A1, A2, and A3. (N) Cuticle of *N.vit mlpt* RNAi larva. Larva is extremely truncated with loss/fusion of all abdominal segments, and fusion of several remaining anterior segments. Only one spiracle remains. (O) Cuticle of *N.vit Svb* RNAi larva. The larva is extremely truncated, with loss of most or all abdominal segments, and fusion of thoracic segments, leaving a single wide band of denticles and one (or two) spiracles in this fused domain. (P) Cuticle of *N.vit ubr3* RNAi larva. Severe phenotypes of *N.vit ubr3* RNAi results in larvae with little or no cuticle that are impossible to recover. Here, the more mild larval phenotype includes a shortened larva with a thin, naked cuticle at the posterior, and few denticles in the anterior.

**Figure 5.**
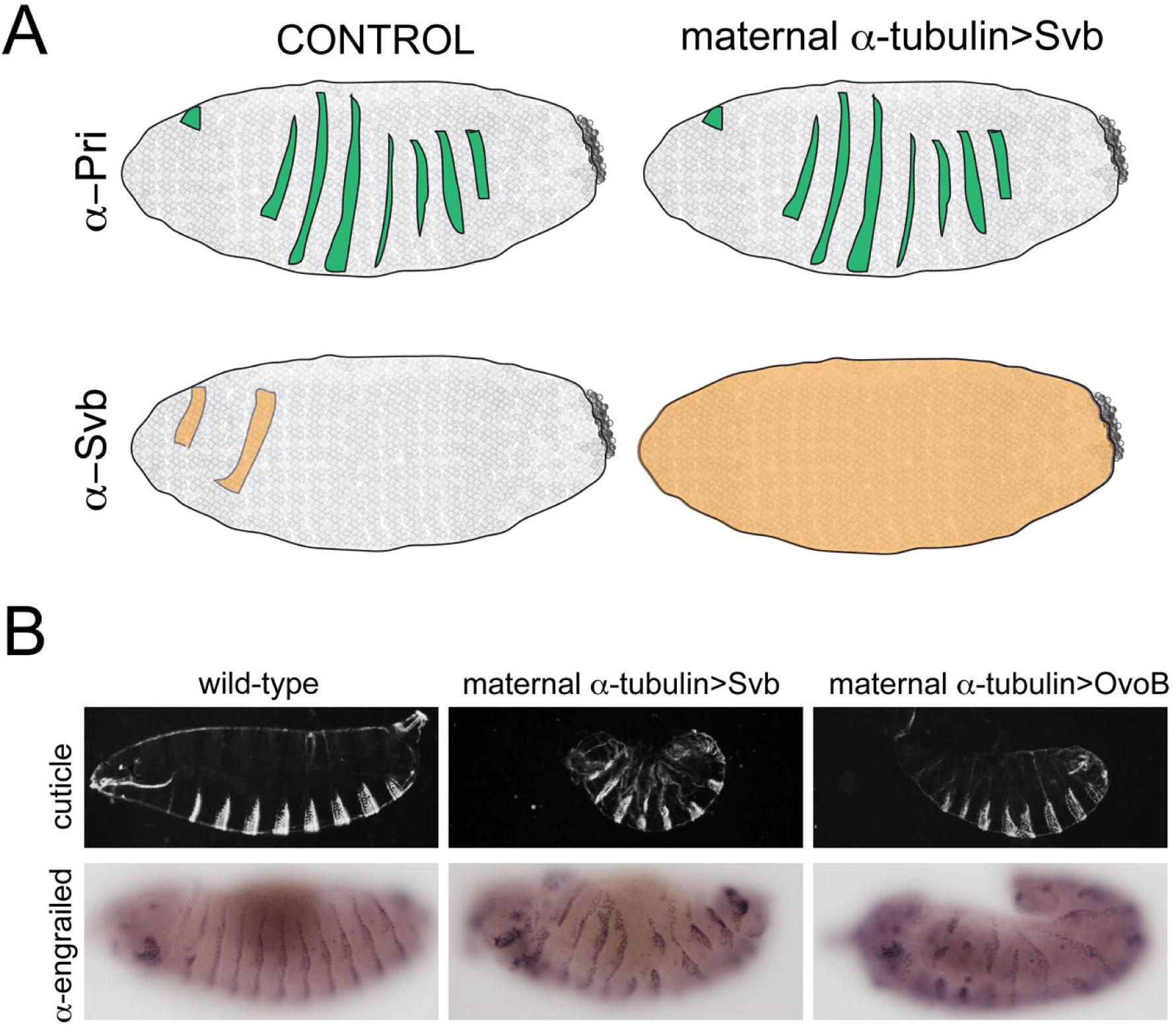
Maternal overexpression of *Dmel Svb* in the early embryo “reawakens” segmentation function. (A) Schematic representation of mRNA expression of both *D. mel pri* and *Svb* in wild type embryos (left embryos) and in embryos expressing *D.mel Svb* maternally from a strong ubiquitous promoter (right embryos). **(B)** Phenotype of maternal expression of *Svb* in *D. melanogaster.* Top panels are larval cuticle preps showing the pattern of ventral denticles that are used as a proxy for proper patterning. Left most panels show denticle pattern and engrailed protein staining in wild type *Drosophila melanogaster.* Middle panels show denticle pattern and engrailed protein staining in *D. mel* overexpressing Svb from a ubiquitous maternal promoter (a-tubulin). Right panels show denticle pattern and engrailed protein staining in *D.mel* overexpressing OvoB activator from the maternal a-tubulin promoter.

### Reawakening functionality *mlpt/svb/ubr3* module in segmentation by restoring Svb expression in the early *Drosophila* embryo

Since we find this functional module to be ancestral and deeply conserved in both short and long germ insects, we sought to determine how the module lost its segmentation role in *Drosophila. ubr3* is ubiquitous and *mlpt* is expressed in pair-rule like stripes, but *svb* expression is absent at these early *Drosophila* embryonic stages. We therefore hypothesized that the loss of the segmentation function of this gene module occurred through the loss of *svb* expression during embryogenesis in the *Drosophila* lineage. To test this hypothesis, we added back *svb* expression to the early embryo context using the *Gal4/UAS* system (Brand AH & Perrimon N, 1993), as it is observed in the blastoderm embryos of *Tribolium, Oncopeltus, Gerris,* and *Nasonia*. Strikingly, over-expression of *svb* in the early embryo resulted in segmentation defects (Figure 5B, middle panels) while loss of *mlpt* function did not (Galindo MI et al, 2007). This result suggests that the function of the *mlpt/svb/ubr3* module in segmentation is contingent upon expression of all three partners and that the loss of this function in flies evolved through the loss of *svb* expression and the dissociation of the trio during early embryogenesis. We next tested whether the segmentation defects observed were due to the processing of the ectopically expressed *svb* into an activator by *mlpt* and *ubr3*. To do this, we over-expressed OvoB, a transcriptional activator originating from the same locus as Svb that behaves like the cleaved form of Svb but that does not require cleavage by Mlpt and Ubr3. Maternal (ectopic) expression of OvoB activator causes segmentation defects that are reminiscent of those obtained by Svb over-expression (Figure 5B, right panels). This result indicates that the segmentation defects observed upon over-expression of either Svb or OvoB are the result of the accumulation of the activator form of Svb/Ovo protein, and that, in this context, ectopic Svb can be regulated and converted to activator form by Mlpt peptides. We therefore conclude that the cooperativity of this gene module has remained intact throughout evolution, and the inactivation of its function in *Drosophila* segmentation likely occurred through the promoter-mediated abrogation of early expression of Svb, an essential component of the module.

## Discussion

In summary, our experiments reveal how a cooperative trio of genes, initially discovered in *Drosophila* where it has a more restricted functional capacity, possesses important ancestral functions in insect embryonic segmentation, representing a significant addition to the anterior-posterior patterning network in insects. Segmentation function of this module has receded in the *Drosophila* lineage through the dissipation of temporal coexpression, while retaining cooperative potency, a solution that preserves connectivity for use in other processes in the fly, such as leg patterning, trichome patterning, and metamorphosis. We surmise that the loss of the essential function of Svb/Mlpt/Ubr3 in segmentation of flies was facilitated by the shift to complete segment patterning before cellularization, which leaves no segments aside for delayed patterning, thus obviating the need for a switch between modes of segmentation. Our current study also illustrates how the use of non-Drosophilid insects with varied modes of embryogenesis can uncover a deeply conserved function, which would have been missed using only the well-studied *Drosophila* model.

We observe several conserved functions of this module of *mlpt, svb* and *ubr3*. All insects from the most basal studied (*Gerris*) to the most derived (*Drosophila*) exhibit strong epidermal patterning defects, evident both in the patterns of apical projections on the larval cuticles of flies and wasps, and in the thinning of the cuticle (Figures 1, 4; Figure 1, Supplement 3) as well as defects in leg patterning in all species examined (Figure 1G, Figure 3). The conserved outputs of this module highlight transcriptional networks downstream of Svb whose connectivity are largely intact over large evolutionary distance, as are sequence features within Svb protein (conserved N-terminus, proteolytic cleavage sites) required for mlpt/ubr3 mediated cleavage (Figure 1-supplement 1). Several *Svb* epidermal targets identified in flies are similarly regulated in *Tribolium* (Li C et al, 2016). Expression patterns of several Svb epidermal target genes in *Nasonia* support this conclusion (Figure 1, Supplement 4). Together, these observations underscore the ancestral conservation of both Svb transcriptional networks and phenotypes dating to early in the radiation of arthropods.

Outside of *Drosophila*, we observe conserved function in insect embryonic segmentation, affecting formation of posterior segments in all insects tested, as well as additional defects in head and thoracic segment derivatives, delineating a new module for insect embryonic segmentation. Our experimental data show that promoter control of Svb expression timing may be sufficient to bring segment patterning potency on-or off-line in the embryo (Figure 5). The phylogenetic distribution within insects of short/long germ modes of development, whose hallmark is delayed abdominal segment formation and patterning, suggests that evolution has likely repeatedly sampled these modes (Misof, B. et al, 2014). Our data suggest one mechanism by which delayed posterior segment formation may be switched on/off via Svb/Mlpt/Ubr3.

The highly dynamic expression of *mlpt* and *svb* and the complementarity of their expression patterns, seen in every non-Drosophilid insect examined in this study, are striking. *Mlpt/svb* boundaries occur at varying positions in different insect embryos. Their expression domains also overlap at times, as in the head in the early *Drosophila* embryo, and later, in the same denticle prepattern. We often observe a sharp boundary in non-*Drosophilid* insects between two domains of strong expression, suggesting mutual exclusivity between high *svb* and *mlpt* expression, or a negative feedback loop. Indeed, we observe derepression of *svb* when *mlpt* is reduced by RNAi (Supplementary Figure 7). The position of the complementary Svb/Mlpt expression boundaries at the interface between patterned segments and unpatterned cells in the growth zone is intriguing. A strong domain of *svb* expression in the growth zone is observed in all short germ species examined, often adjacent to a strong *mlpt* expression domain. In cells distant from the boundary, high levels of Svb repressor protein that excludes Mlpt expression would silence target promoters, while Svb targets in the region of overlap at the junctions between the *svb*-*mlpt* domains would be activated due to more moderate Svb repressor levels that are no longer inhibitory to Mlpt expression. Alternatively, potential non-cell autonomous activity of Mlpt peptides could activate Svb and its target promoters close to the boundary even in the absence of an overlap between the mRNA expression domains. The potential of this interface to produce a narrow adjacent domain of Svb target activation is indeed interesting. The short-range gradient of Svb activator could constitute a retracting wavefront (regulated by or acting in conjunction with a speed/frequency regulator; Zhu, X et al, 2017) that determines spatial patterning. The position of the complementary Svb/Mlpt expression boundaries at the interface between blastoderm and oscillation-driven growth zone invites further study.

Beyond insects, Svb/Ovo is conserved in all animals, and predates bilateria (Kumar A et al, 2012). Svb in animals functions “classically” in germline and epidermis, though evidence now suggests epithelial function in mouse, as well (Dai, X. et al, 1998; Nair, M. et al. 2006). Svb mammalian homologs are involved in phenotypic changes in cells, including significant cytoskeletal rearrangements, and mesenchymal to epithelial transitions (MET; Ito, T. et al, 2017; Roca, H. et al., 2013; Zanet J, et al., 2015; 28. Jia, D. et al, 2015). Insect gastrulation, which also relies heavily on cytoskeletal rearrangements during germband elongation and germ layer growth (Collinet, C. et al, 2015; Munjal, A. et al, 2015) may share this feature, since embryos with reduced Mlpt or Svb appear deficient in convergent extension and directional growth, implicating cytoskeletal effectors. Importantly, both in mammalian models and invertebrates, the repressor form of Svb/Ovo transcription factor has different activity than the activator form (Masu, Y. et al, 1998; Hayashi, M. et al, 2017). Insects achieve switching between these activities through post-translational conversion of Svb protein through proteolytic processing, while vertebrate Ovo proteins possess different promoters controlling transcript variants with divergent transcriptional activity (Li, B. et al, 2002; Masu, Y. et al, 1998). Together, our data suggest how a post-translational mechanism involving a micropeptide like Mlpt can be used to switch Svb activity, broadly regulating phenotypic plasticity during embryogenesis.

In conclusion, our experiments build on the strength of deep evolutionary understanding of a micropeptide switch and its genetic context to revive ancestral functionality in an essential developmental network. This process demonstrates potential for the rational design of novel biological switches, using evolutionarily conserved components in contexts in which they no longer possess native function, but in which they are poised to do so.

## Materials and Methods

### Experimental Design

All insect cultures were reared at ambient temperature of 25^°^C, unless otherwise noted. Wild type ***<u>Nasonia</u>*** embryos were collected from virgin AsymCx (*34*) females host fed on *Sarcophaga bullata* pupae (Carolina Biological), aged as needed at 25^°^C, and fixed for 25 minutes in 4% heptane-saturated formaldehyde/1X Phosphate Buffered Saline (PBS), with vigorous shaking. Embryos were transferred to 100% methanol for storage, or further processed. Embryos were then rehydrated to 1X PBS with 0.1% Tween (PBT) through a methanol/PBT series, and hand peeled under 1X PBT using 1mL insulin needles (Becton-Dickinson).

Wild-type *<u>Oncopeltus</u>* embryos were collected on cotton from mated females, and aged, as needed, in a 25^°^C incubator. Embryos were first boiled for 1 to 3 minutes in a microfuge tube in water, followed by a 1 minute incubation on ice, before further processing. Embryos were fixed in 12% heptane-saturated formaldehyde/1X PBS for 20 minutes with shaking. The heptane was replaced by methanol, and the embryos either stored under methanol at −20^°^C or processed immediately. Embryos were then rehydrated to 1X PBT through a methanol/PBT series, and dechorionated, before further fixation for 90-120 minutes in 4% formaldehyde/1X PBT. Embryos were then transferred to and stored in 100% methanol.

Wild type ***<u>Gerris</u>*** *buenoi* were collected from a pond in Toronto, Ontario, Canada and established in the lab. Stocks were maintained in aquaria at 25 ^°^C with a 14-h light/10-h dark cycle, and fed with fresh crickets. Styrofoam float pads were provided to females as substrate for egg lays. Embryos were collected and incubated at 20-25^°^C until desired developmental timepoints, at which time they were dissected in 1x PBS with 0.05% tween-20 (“PTW”). Once dissected, embryos were fixed in 4% paraformaldehyde and stored under 100% Methanol at −20^°^C until use.

Wild type ***<u>Drosophila</u>*** lines were reared at ambient temperature under standard conditions.

Wild type ***<u>Tribolium</u>*** embryos were fixed and single whole-mount in situ hybridizations were performed as previously described (*35, 36, 37*). Digoxigenin-labeled RNA probes were detected using alkaline phosphatase-conjugated anti-DIG antibodies (1:2000; Roche) and NBT/BCIP substrates (Roche) as per manufacturer instructions.

#### Antibody staining of Tribolium embryos

Freshly fixed embryos were washed with PBT (PBS and 0.1% Triton X) and exposed to rabbit anti-Svb 1S at a concentration of 1:2000 for 4h at room temperature on a rotating wheel. Following washes, the embryos were incubated with preabsorbed alkaline phosphatase conjugated goat anti-rabbit secondary antibody at a concentration of 1:1000 for 2h at room temperature on the wheel. Following washes the embryos were stained with NBT-BCIP substrate and visualized under a stereomicroscope.

### Parental RNAi (pRNAi) and larval/hatchling cuticle preparation

#### Tribolium pRNAi

dsRNA synthesis and parental RNAi were performed as described previously (*38*; *39*). dsRNAs were injected into female pupae or virgin adult females at a concentration of 1-3μg/μl. RNAi phenotypes were confirmed by injection of non-overlapping dsRNA fragments.

First instar larval cuticles were cleared in Hoyer’s medium/lactic acid (1:1) overnight at 60^°^C. Cuticular autofluorescence was detected on a Zeiss Axiophot microscope. Z stacks and projections were created with a Zeiss ApoTome microscope using the Axiovision 4.6.3.SP1 Software. Colour images were taken by (ProgResC14) using the ProgResC141.7.3 software and maximum projection images were created from z stacks using the Analysis D software (Olympus).

#### Oncopeltus pRNAi

dsRNA template was amplified from target gene fragments which had been cloned into either pCR-Topo (Qiagen) or pGEM (invitrogen), using T7 promoter-containing oligos, as described previously (*40*). Purified PCR product was used for dsRNA transcription using *Megascript RNAi* (Ambion) according to manufacturer’s instructions. dsRNA was injected into newly eclosed virgin female milkweed bugs, at a concentration of 1-3μg/μl. After injection, females were mated to uninjected males, and embryos were collected for the duration of egglaying. Embryos for phenotypic evalution were incubated at 28^°^C for 8 days, and unhatched embryos were dissected from their membranes and imaged for hatchling phenotypes.

#### Gerris buenoi parental RNAi

Knock down of *pri, svb* and *ubr3* genes was performed by injection of double-stranded RNAi into adult females abdomen (*41*). dsRNA was synthesized by *in vitro* transcription and purified following the protocol described in (*40*) Nature Protocols. dsRNA was injected at a concentration of 1ug/uL in 1X injection buffer prepared following the protocol in (*43*). dsRNA directed against the *yellow fluorescent protein* gene was used as an injection control.

#### Nasonia embryo RNAi

dsRNA template was amplified from target gene fragments that had been previously cloned into pCR-Topo (Qiagen) or directly from embryo cDNA, using standard T7 promoter-containing oligos, as described previously (*41*). Purified PCR product was used for dsRNA transcription using *Megascript RNAi* (Ambion) according to manufacturer’s instructions, and purified product diluted to 1-3μg/μl for injection. pRNAi for *mlpt* and *svb* in *Nasonia* resulted in sterility. Therefore, haploid embryos laid by unmated host-fed virgin *Nasonia* females were microinjected with dsRNA using a Femto-Jet microinjector (Eppendorf), and transferred to a slide to develop in a humid chamber at 28^°^C for 36H. Unhatched larvae were dissected from extraembryonic membranes and cleared in freshly prepared Lacto:Hoyer’s medium overnight at 65^°^C, and imaged for cuticle autofluorescence the following day.

#### Drosophila UAS/Gal4 experiments

The following Drosophila lines were used in this study: w, pri1/TM6, Ubi-GFP, svbR9/ FM7, Kr-GFP, nullo-Gal4 (from Gehring lab), mat-Gal4, nanos-Gal (gift from N. Dostatni). UAS drivers used in this study are as follows: UAS-svb, UAS-ovoA, UAS-ovoB, UAS-pri. Ubr3 mutant embryos devoid of maternal and zygotic contribution were generated using the Ubr3B allele according to (*15*).

### *In situ* hybridizations and immunostaining

#### Gerris buenoi

*In situ* hybridizations in *Gerris* were performed as previously described (*41*). Briefly, embryos were rehydrated to 1X Gerris PBT, through a MeOH/PTW series, and then washed 3 times in PTW to eliminate residual methanol. Embryos were then permeabilized in PBT 0.3% and PBT 1% (1X PBS; 0.3% or 1% Triton X100) for 1 hour. Following these washes, embryos were rinsed once for 10 minutes in a 1:1 mixture of PBT 1% and Hybe B (Hybe B is: 50% Formamide; 5% dextran sulfate; 100 mg/ml yeast tRNA; 10X salts). The 10X salt mix contains 3 M NaCl; 100 mM Trizma Base; 60 mM NaH2PO4; 50 mM Na2HPO4; 5 mM Ficoll; 50 mM PVP; and 50 mM EDTA. RNA probes corresponding to each gene were transcribed from cDNA templates cloned into pGEM-T (Promega), and then transcribed *in vitro* using either T7 or Sp6 RNA polymerase (Roche) and labeled with Digoxigenin-RNA labeling mix (Roche). Pre-incubation of embryos was carried out in Hybe-B for 1 hour at 60^°^C before adding Digoxigenin-labeled RNA probes overnight at 60^°^C. The next day, embryos were washed in decreasing concentrations of Hybe-B diluted with PBT 0.3% (with 3:1, 1:1, 1:3) and then rinsed three times 5 minutes each in PBT 0.3% and then once for 20 minutes in blocking solution (1X PBS; 1% Triton X100; 1% BSA) at room temperature before adding alkaline phosphatase conjugated anti-DIG antibody (Roche). Embryos were incubated with primary antibody for 2 hours at room temperature. Following primary antibody, embryos were washed for 5 minutes in PBT 0.3% and and then 5 min in PTW 0.05%. Color enzymatic reaction was carried out using NBT/BCIP substrate (Roche) in alkaline phosphatase buffer (0.1M Tris pH 9.5; 0.05M MgCl2; 0.1M NaCl; 0.1% Tween-20), according to manufacturer’s instructions. Upon completion, the reaction was stopped with several washes of PBT 0.3% and PTW 0.1% (1xPBS; 0.1% Tween-20). Stained embryos were stored in 50% Glycerol/1x PBS at 4^°^C or −20^°^C until mounting on slides in 80% glycerol for imaging.

### Immunostaining on *Gerris* embryos

Embryos were cleaned with dilute bleach solution and washed PTW 0.05%. After dissection, embryos were fixed for 20min in 4% Formaldehyde/1X PTW 0.05%. Embryos were then permeabilized with PBT 0.3% for 30 minutes and incubated in antibody blocking solution (1X PBS; 0.1% Triton X100; 0.1% BSA; 10% NGS) at room temperature for 1 hour. Embryos were transferred to blocking solution containing primary antibody (mouse anti-Ubx-AbdA Hybridoma bank 1:5 dilution; rabbit anti-Dll (*42*), catalog number; 1:200 dilution) and incubated overnight at 4^°^C. The next day embryos were washed in PTW 0.05% (two quick washes, then two longer washes 10 min each) and incubated for 30 minutes in blocking solution at room temperature with shaking, before adding the secondary antibody (Rabbit anti-mouse-HRP [1:1000] from Promega or Donkey anti-Rabbit-HRP [1:500] from Jackson Immuno research) diluted in PTW. All secondary antibodies were incubated with embryos for 2 hours at room temperature with shaking. Following secondary antibody, embryos were rinsed in PBT 0.3% and PTW 0.05% three times each for 10 minutes at room temperature. Before enzymatic developing with DAB with color enhancer (Diaminobenzidine tetrahydro-Chloride from Sigma), embryos were briefly incubated with DAB solution for 5 minutes at room temperature. Upon completion, staining was stopped by washing the embryos briefly in PBT 0.3%, followed by 5 times, five minute washes in PBT 0.3%. Five more washes of 5 minutes in PTW 0.1% followed. Embryos were transferred to 30% glycerol/1X PBS for 5 min, and then 50% Glycerol/1X PBS for 5 min, before sinking in 80% glycerol/1X PBS at 4^°^C until mounting in 80% glycerol under coverslips for imaging.

### *Oncopeltus in situ* hybridizations

were carried out (as described for *Nasonia* in *14)* on embryos peeled and stored under 100% methanol, and rehydrated through an methanol/1x PBS, 0.1%Tween (1xPBT) series. Briefly, rehydrated embryos were washed several times in 1x PBT before a 5 minute post-fix in 5% formadelhyde/1X PBT, followed by 3 five minute washes in 1X PBT. Embryos were briefly treated with Proteinase K (4μg/mL final concentration) in 1X PBT for 5 min, followed by 3 five minute washes in 1X PBT, and an additional 5 minute post-fix in 5% formaldehyde/1X PBT. Following 3 x three minute washes in 1X PBT, embryos were incubated in hybridization buffer for 5 minutes at room temperature, followed by incubation in fresh hybe buffer for a 1 hour pre-hybridization step at 65^°^C. RNA probes were prepared and added to a fresh portion of hybridization buffer and incubated at 85^°^C for 5 minutes, then one minute on ice, before replacing pre-hybe with hybe buffer containing denatured RNA probe. Tubes were incubated overnight at 65^°^C. After washes in formamide wash buffer, embryos were washed in several changes of 1X MABT buffer, before incubation in 1X MABT+ 2% Blocking Reagent (BBR; Roche Diagnostics) for 1 hour, and then 1X MABT/2%BBR/20% sheep serum for an additional hour, before addition of fresh 1X MABT/2%BBR/20% sheep serum containing anti-DIG AP Fab fragments (1:2000; Roche) for overnight incubation at 4^°^C. In the morning, extenstive 1X MABT washes were carried out before equilibration of embryos with AP staining buffer and then staining with AP staining buffer containing NBT/BCIP (Roche; as per manufacturers instructions). After staining, three 1X PBT washes were carried out before a final post-fixation step (5% formaldehyde/1xPBT), and then one PBT wash before sinking in 50% glycerol/1xPBS, and then 70% glycerol/1xPBS, which was also used for mounting before imaging.

#### Nasonia in situ hybridizations

*Nasonia* embryos were collected and fixed in 4% formaldehyde/1X PBS/Heptane for 28 minutes with moderate shaking. The embryos were hand-peeled under 1x PBS+ 0.1% tween, after being affixed in room air to double sided tape, and hand-peeled as described previously, except that embryos were collected from host-fed, mated females. Fixed embryos were stored under methanol at −20^°^C until use.

*In situ* hybridizations were carried out as described previously (*44).* Briefly, fixed embryos that had been stored under methanol were gradually brought up to 1X PBT in a PBT/MeOH series, and washed three times in 1x PBS +0.1% tween-20 (PBT) before a 30 minute post-fixation in 5% formaldehyde/1XPBT. The embryos were then washed three times in 1X PBT, and digested in Proteinase K for five minutes (final concentration of 4mg/mL) before three PBT washes. Embryos were blocked for 1 hour in hybridization buffer before probe preparation (85^°^C, 5’; ice 1’) and addition for overnight incubation at 65^°^C. The next day, embryos were washed in formamide wash buffer three times, and and then 1X MABT buffer three times, before blocking in 2% Blocking Reagent (BBR; Roche Diagnostics) in 1X MABT for 1 hour, then in 10% horse serum/2% BBR/1XMABT for 2 hours. Embryos were incubated overnight at 4^°^C with primary antibody (anti-DIG-AP Fab fragments; Roche, 1:2000). The third day, embryos were washed in 1X MABT for ten x 20 minutes washes before equilibrating the embryos in AP staining buffer and developing in AP buffer with NBT/BCIP solution (Roche Diagnostics). After staining, embryos were washed in 1x PBT three times for five minutes each before a single 25 minutes post-fix step in 5% formaldehyde/1XPBT. Embryos were then washed several times with 1X PBT, and allowed to sink in 50% glycerol/1XPBS and then 70% glycerol/1XPBS, which was subsequently used for mounting.

#### Cloning of Tribolium cDNA sequences

For Tribolium svb, all primer pairs shown were used to generate template for dsRNA synthesis. Amplicons generated by the last four pairs were also used for antisense RNA probe synthesis. dsRNA fragments corresponding to different regions of the svb transcript were used for gene knockdown by RNAi. All dsRNA fragments resulted in similar knockdown phenotypes with high penetrance. Primers were designed based on the Next-RNAi software, Primer3 or MacVector. The nucleotides shown in red indicate tags of parts of T7 (3’ primer) and SP6 (5’ primer) promoter sequences attached to gene-specific sequences in the manner described by Schmitt-Engel et al. 2015. The products were used for a second PCR using T7 and SP6-T7 primers for generating a double stranded template for in-vitro transcription by T7 polymerase. For in-situ RNA probes, the second PCR was done using the complete T7 and SP6 promoter sequences and subsequently in-vitro transcription was performed to generate a digoxigenin-labeled antisense RNA probe with the appropriate polymerase. Amplicons that were cloned into pBluescript vector were amplified with T7 and T3-T7 primers for subsequent dsRNA synthesis or T7 and T3 primers for subsequent antisense RNA probe synthesis using either T3 or T7 RNA polymerase. The primer design was based on the RNAseq data (Tcas au5 prediction) for svb available on iBeetle-Base. For mlpt dsRNA and probe synthesis, a full-length mlpt cDNA cloned into pBluescript was obtained from Dr. Michael Schoppmeier. For ubr3, all primer pairs shown were used for dsRNA synthesis. All dsRNA fragments resulted in similarly strong knockdown phenotypes with very high penetrance. The fragments generated with the primers containing iBeetle numbers were also used as probes.

#### Generating a Tcas svb mutant using CRISPR/Cas9

The *svb* gRNAs were directed to the putative transactivation domain in exon 2 of *Tc-svb*. The G (required by the U6 promoter for transcription initiation) is marked in green. The PAM sequence is shown in blue. The sequences in orange represent the complementary overhangs generated by *Bsa1* digestion. A fourth gRNA was directed to the *Tribolium single-minded* gene (*Tc-sim*, Rode and Klingler, unpublished). Embryonic injection mix consisted of 125 ng each of the four gRNA expression vectors, 500ng of the donor eGFP vector containing the *sim* target sequence, and 500ng of the Cas9 expression vector. Non-homologous end joining (NHEJ) method was employed for directed knock-in of an eGFP containing donor marker plasmid into the exon2 of the endogenous *svb* gene. The *sim* gRNA was used to target the *sim* sequence in the marker plasmid leading to its Cas9-induced linearization. This was followed by insertion of the linearized plasmid into one or more target sites in the *svb* genome. A successful knock-in of the marker plasmid was obtained only at gRNA target site 3. This insertion site was present in all *svb* transcripts and was also downstream from a putative second start codon, thus increasing the chances of obtaining a *svb* null phenotype.

## Acknowledgments

MIR would like to thank Ariel Chipman, Claude Desplan, and Igor Ulitksy for scientific support during the course of this work and for critical reading of the manuscript.

## Author contribution

Projects were conceptualized and supervised by SR, MIR, HC-D, AK, MK and FP. The original draft of the manuscript was written by SR and MIR, and subsequently revised mainly by SR, MIR, HC-D, AK, MK and FP (all authors edited and read the final version). Experiments were planned and conducted by SR, MIR, HC-D, AD, BS, WT, TA, IS, MT and FP.

## Data and materials availability

Sequences presented in this paper can be found in Genbank, with accession numbers as follows: *Tcas Svb* MG913606 *Nvit mlpt* MH181829 *Nvit Svb* MH181831 *Nvit Ubr3* MH181828 *Ofas mlpt* MH181830 *Ofas svb* MH181832 *Ofas Ubr3* MH 181827 *Gbue Svb* MH011417 *Gbue mlpt* MH011418 *Gbue Ubr3* MH011418.

